# Spectral and topological analysis of the cortical representation of the head position: does hypnotizability matter?

**DOI:** 10.1101/442053

**Authors:** Esther Ibáñez-Marcelo, Lisa Campioni, Diego Manzoni, Enrica L. Santarcangelo, Giovanni Petri

## Abstract

The aim of the study was to assess the EEG correlates of head positions, which have never been studied in humans, in participants with different psychophysiological characteristics, as encoded by their hypnotizability scores. This choice is motivated by earlier studies suggesting different processing of the vestibular/neck proprioceptive information in subjects with high (*highs*) and low (*lows*) hypnotizability scores maintaining their head rotated toward one side (RH). We analysed EEG signals recorded in 20 highs and 19 lows in basal conditions (head forward) and during RH, using spectral analysis, which captures changes localized to specific recording sites, and Topological Data Analysis (TDA), which instead describes large-scale differences in processing and representing sensorimotor information. Spectral analysis revealed significant differences related to the head position for alpha1, beta2, beta3, gamma bands, but not to hypnotizability. TDA instead revealed global hypnotizability-related differences in the strengths of the correlations among recording sites during RH. Significant changes were observed in *lows* on the left parieto-occipital side and in *highs* in right fronto-parietal region. Significant differences between the two groups were found in the occipital region, where changes were larger in *lows* than in *highs*. The study reports findings of the EEG correlates of the head posture for the first time, indicates that hypnotizability modulates its representation/processing on large-scale and that spectral and topological data analysis provide complementary results.

## 1 Introduction

The EEG correlates of the tonic vestibular/neck proprioceptive information related to head positions have never been studied in humans. It has been reported, however, that the vestibular and neck proprioceptive information are conveyed to fronto-temporo-parietal cortex, insula and hippocampus [De Waele et al., 2001, Lopez and Blanke, 2011], that galvanic vestibular stimulation induces a slight suppression of gamma power in lateral regions followed by an increase in beta and gamma power in frontal regions, and that the power of each oscillatory band throughout frontal, central/parietal and occipital electrodes are linearly correlated with the stimulus intensity [Kim et al., 2013].

Recent evidence has shown that sensori-motor integration is modulated by the psychophysiological trait of hypnotizability [Santarcangelo and Scattina, 2016]. Hypnotizability is known to predict the proneness to modify perception, memory and behavior according to specific suggestions and is measured by scales. Alterations of visual and leg proprioceptive information [Santarcangelo et al., 2008a] and asymmetric tactile feet stimulation [Solari et al., 2016] induce larger and/or faster body sway in highly hypnotizable individuals (highs), while tonic neck rotation induces changes in the velocity of body sway only in low hypnotizable subjects (lows) [Santarcangelo et al., 2008b]. Hypnotizability is also associated with morpho-functional differences in the cerebral [Landry et al., 2017] and cerebellar cortex ([Picerni et al., 2018, Bocci et al., 2017], which are structures relevant to sensori-motor integration. Indeed, the vestibular and neck proprioceptive informations are conveyed to fronto-parietal, insular and cingulate cortices which show also hypnotizability-related morpho-functional properties [Landry et al., 2017]. In addition, the cerebellar function, which is modulated by hypnotizability [Bocci et al., 2017], is required for the elaboration of sensori-motor information related to the head position [Manzoni, 2005, Kammermeier et al., 2009]. Spectral analysis of EEG signals can be used to characterize the cortical representation of the head posture. In fact, beta power is reduced in all conditions related to movement actual, observed and imagined movement/posture [Pfurtscheller et al., 2005, Keinrath et al., 2006, Turella et al., 2016], whereas it may increase during action planning without execution, which is likely due to the integration of bottom-up sensorimotor information due to movement [Turella et al., 2016]. Gamma power synchronization has been found in a wide range of cognitive [Bosman et al., 2014] and sensory operations [van Ede et al., 2014]. Alpha power is mainly involved in cognitive processes [Başar et al., 1997, Klimesch, 1999] but modulation of alpha rhythms has been observed also during the elaboration of event-specific sensory and motor information [Babiloni et al., 2016] and is influenced by the specific task and by the individual motor experience [Duru AD, 2018]. Hypnotizability-related EEG spectral differences have been studied during imagery tasks [Cavallaro et al., 2010], but not during different sensori-motor conditions, with the exception of the nociceptive stimulation [Zeev-Wolf et al., 2016] which, however, has been preferentially investigated through cortically evoked potentials [De Pascalis et al., 2015, Valentini et al., 2013].

Spectral observables characterize local changes in the cortical activity, whereas topological ones, obtained from Topological Data Analysis techniques (TDA, Appendix 3.2), are able to characterize the shape and properties of the networks active in correspondence of specific conditions on a mesoscopic scale [Sporns, 2013, Petri et al., 2014, Lord et al., 2016, Giusti et al., 2016, Sizemore et al., 2018]. Specifically, TDA was instrumental in highlighting different global processing schemes of the vestibular/neck proprioceptive information related to the rotated position of the head (RH) in highs and lows. In fact, during both real and imagined rotated head position the former display smaller changes than the latter with respect to basal, head forward conditions [Ibanez-Marcelo et al., 2018]. The highs EEG characteristics are likely due to a moderate but widely distributed restructuring of the brain activity, rather than to the occurrence of pronounced local changes during RH. Previous TDA analyses investigated global and mesoscopic properties on the active networks [Ibanez-Marcelo et al., 2018], rather than possible spatial differences between conditions and groups. We take here an alternative approach by focusing on the comparison of both classical spectral analysis and the study of topological properties at the nodal level, that is, with reference to each recording site. Persistent homology (Appendix 3.2), one of the main tools of TDA, describes the shape of high-dimensional datasets by producing a series of progressively finer approximations of a given whole data space. It studies the evolution of the connectivity and lack thereof, hence holes, in all dimensions (e.g. connected components, one-dimensional cycles, three-dimensional cavities, and their higher dimensional analogues [Sizemore et al., 2016] along this sequence of approximations (called a filtration). Here, the data space we focus on is the one generated by the correlation matrices between EEG signals of each subject. The persistent homology of these spaces captures the both the presences and lack of correlation patterns between multiple recording sites. Cycles, each corresponding to a region of weakened connectivity in the correlations patterns, are mesoscopic topological features which encompass multiple regions and are characterized by their appearance along the series of approximation (cycle birth, b), their disappearance (death d), the number of regions they include (cycle length) and their persistence across the filtration, defined as pi=d-b, capturing how long they live. Local topological information about specific regions can be obtained from the homological scaffold [Petri et al., 2014]: a network representation, where of the homological structure built by aggregating the cycles and weighting them according to their persistence. This network is composed by nodes, which represent recording sites, and edges, that represent connections between such sites. The intensity of the connection is represented by a weight assigned to the edge. A measure of the importance of a node, which we dub nodal strength, is then obtained by summing the weights of the edges stemming from that node in the scaffold. This method has been applied to resting state fMRI data and has revealed topological correlates of altered states of consciousness [Petri et al., 2014] and epileptic seizures [Santarcangelo and Scattina, 2015], as well as pointing to specific topological structures in resting state [Lord et al., 2016] and during attention modulation [Yoo et al., 2016]. The aim of the present exploratory study was to assess the electroencephalographic correlates of the position of rotated head in highs and lows as reflected by spectral properties of the alpha, beta and gamma frequencies and by the topological structure of the correlations among EEG signals.

## 2 Methods

### 2.1 Subjects

The study was approved by the local Ethics Committee. After signing an informed consent, hypnotizability was measured through the Italian version of the Stanford Scale of Hypnotic Susceptibility (SSHS), form A [Weitzenhoffer and Hilgard,] in a sample of 200 right-handed (Edinburgh Handedness Inventory) students of the University of Pisa with negative anamnesis of neurological and psychiatric disease and drug free for at least the latest 2 weeks. They were classified as highly (highs, SHSS score ≥ 8/12), medium (mediums, SHSS score: 57) and low hypnotizable subjects (lows, SHSS score < 4/12). Twenty consecutive highs (SHSS score (mean+SD): 9.6 ± 1.4, 11 females) and 20 consecutive lows (SHSS, mean+SD: 1.5 ± 0.9, 11 females) were enrolled.

### 2.2 Experimental Procedure

Experimental sessions were conducted between 11a.m and 2 p.m., at least 2 h after the last meal and caffeine containing beverage. Participants were comfortably seated in a semi-reclined armchair in a sound-and light-attenuated room. After EEG montage, they were invited to close their eyes and relax at their best for 1 minute (basal conditions (B), head forward) and then to rotate their head toward the right side to align their chin with the shoulder and maintain this position for 1 minute (rotated head, RH). The head position and eye closure were visually controlled throughout the session by one of the experimenters.

### 2.3 EEG acquisition and processing

EEG was acquired at a sample rate of 1000 Hz through a Quick-CapEEG and Quick-Cell system (Compumedics NeuroMedical Supplies). EEG electrodes (N=32) were placed according to the1020 International System. Auricular, eye and EKG electrodes (standard DI lead) were also placed. The reference electrode during acquisition was FCz; off-line the signal was referred to A1/A2 and FCz was restored. Filters (notch at 50 Hz, bandpass 0.545 Hz) were applied a-posteriori. Electrodes impedance was kept under 10 *k*Ω. No participant had more than 1 bad channel, interpolated using the spherical interpolation method (EEGLAB pop interp function). Source components were obtained using Indipendent Component Analysis (infomax ICA algorithm, EEGLAB function runica). They were visually inspected to remove artefacts. The signal was divided into 20 s epochs (20.000 samples). No windowing function was applied to the raw data. The epoch selection process (deletion criteria: amplitudes *≥* 100*μV*, median amplitude*>*6SD of the remaining channels) removed a maximum of 1 epoch in each subject. The earliest less noisy 20 s interval of basal conditions and the interval comprised between the 20th and 40th sec of RH were chosen for analysis. Absolute spectral powers were estimated on two separate 10 s epochs using the Welchs method.

### 2.4 Persistent Homology

In the same time intervals topological invariants were extracted from the correlations among EEG signals during basal and task conditions. In particular, persistent homology (see Appendix 3.2 for details) was computed as follows:

- For each subject and condition, we computed the (Pearson) correlation matrix between all pairs of EEG signals (contained in (-1,1)). We then consider a similarity matrix defined as *S_ij_* = 1 *− |C_ij_|*.
- Each matrix was then thresholded at all distinct values between -1 and 1 to produce a sequence of approximated similarity matrices (the first matrix is empty, while the last contains the entire information). To each thresholded matrix, we can associate a new structure, called a simplicial complex, the shape and structure of which depends on the signal properties and can be characterized by the number and properties of its holes; we focus here on one-dimensional cycles.
- Each cycle is characterized by its birth, death, length and persistence across the sequence of thresholded matrices. For each subject, an individual homological scaffold was then constructed by aggregating all the cycles and summing the associated weights. In this way, it is possible to project the persistent homological information to the level of regions and compute the nodal strength, that is, the integrated amount of cycles that pass through each node, capturing its topological importance.
- For each region/node *r*, we constructed a vector 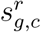, containing the nodal strengths of all subjects in a certain group g (highs, lows) and condition c (basal, RH). The ith entry 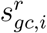 in each vector corresponded to the nodal strength of the ith subject in group g and condition c. We then computed node-specific differences at the group level by measuring the Euclidean distance 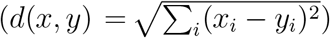, between the vectors corresponding to basal and task condition, where each component *i* corresponds to a subject in the same group. The values obtained measure the extent of the change between basal and condition at the group level for a specific region.

### 2.5 Variables and Statistical Analysis

The log transformed absolute power of beta1 (1316 Hz), beta2 (1620 Hz), beta3 (2036 Hz) and gamma bands (36-45 Hz) recorded during simple relaxation with head forward (basal, B) and in conditions of rotated head (RH) were studied. The absolute power of each frequency band averaged across the left (Fp1, F3, F7) and right frontal (Fp2, F4, F8), left (FC3, FT7, T3) and right medio-anterior (FC4, FT8, T4), left (C3, TP7, CP3) and right medio-posterior (C4, TP8, CP4), left (T5, P3, PO1, O1) and right occipital (T6, P4, PO2, O2) and anterior (Fz, Fcz, Cz) and posterior midline region (CPz, Pz, Oz) was studied. Repeated measures ANOVA (SPSS.15) according to a 2 Hypnotizability (highs, lows) *×* 2 Conditions (B, RH) *×* 2 Sides (right, left) design. The Greenhouse-Geisser ϵ correction for non sphericity was applied when necessary. Post hoc comparisons were performed through paired t-test between conditions and unpaired t test between groups. The same design was applied to the analysis of births, deaths, lengths, persistences of cycles. Since the number of nodes involving more than one hemisphere war much larger than the number of hemispherically restricted nodes, the nodal strengths in the scaffolds was analysed through a 2 Hypnotizability *×* 2 Conditions design. For all analyses the significance level was set at p=.05.

## 3 Results

One low subject was excluded from spectral analysis due to noisy EEG signals, thus the findings were obtained in 20 highs and 19 lows. Persistence homology could be studied in 18 highs and 19 lows because in 2 highs TDA did not detect cycles. The absence of detectable cycles indicate a locally uniform correlations structure, which is in line with the increased uniformity found among highs [Ibanez-Marcelo et al., 2018].

### 3.1 Spectral Analysis

Spectral analysis revealed significant differences between head positions, but not between highs and lows. As reported in Table C.1, ANOVA revealed that in the frontal and medio-anterior regions all frequency bands exhibited significant Conditions effects indicating that their absolute power increased during the maintenance of the rotated position of the head (RH) with respect to basal conditions (B). In the medio-posterior and occipital regions significant increases were observed for beta2, beta3 and gamma. Significant Condition *×* Side interactions (Table C.1) were observed at frontal level for beta3 and gamma, at medio-anterior level for beta2, beta3 and gamma (Figure 1A-C) and at medio-posterior level for gamma. In all these regions, during RH the EEG bands power increased on both sides, but the increases were higher on the right than on the left side. In contrast, at medio-posterior level a significant decrease in the alpha1 power was observed during RH on the right side and no significant change was found on the left side (Figure 1D). On the midline, significant increases in beta2, beta3 and gamma power were observed during RH (Table C.1).

**Figure 1:**
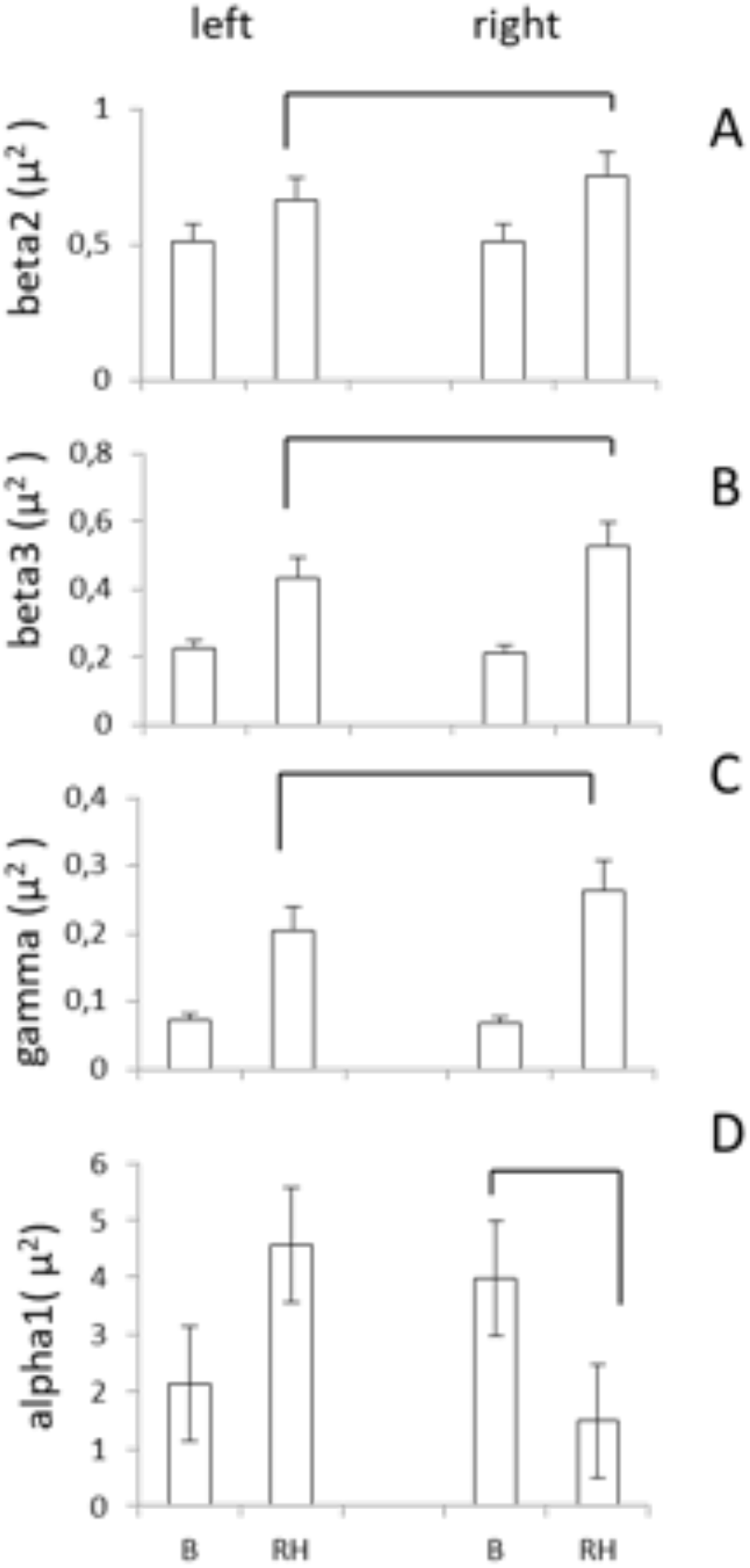
Spectral results. Original absolute power (mean, SEM), collapsed left and right frontal sites. B, basal, head forward condition; RH, rotated head. Lines indicate significant differences. Filled dots are sites showing significant differences in the nodal strength distribution (red *highs*, blue *lows*)

**Figure 2:**
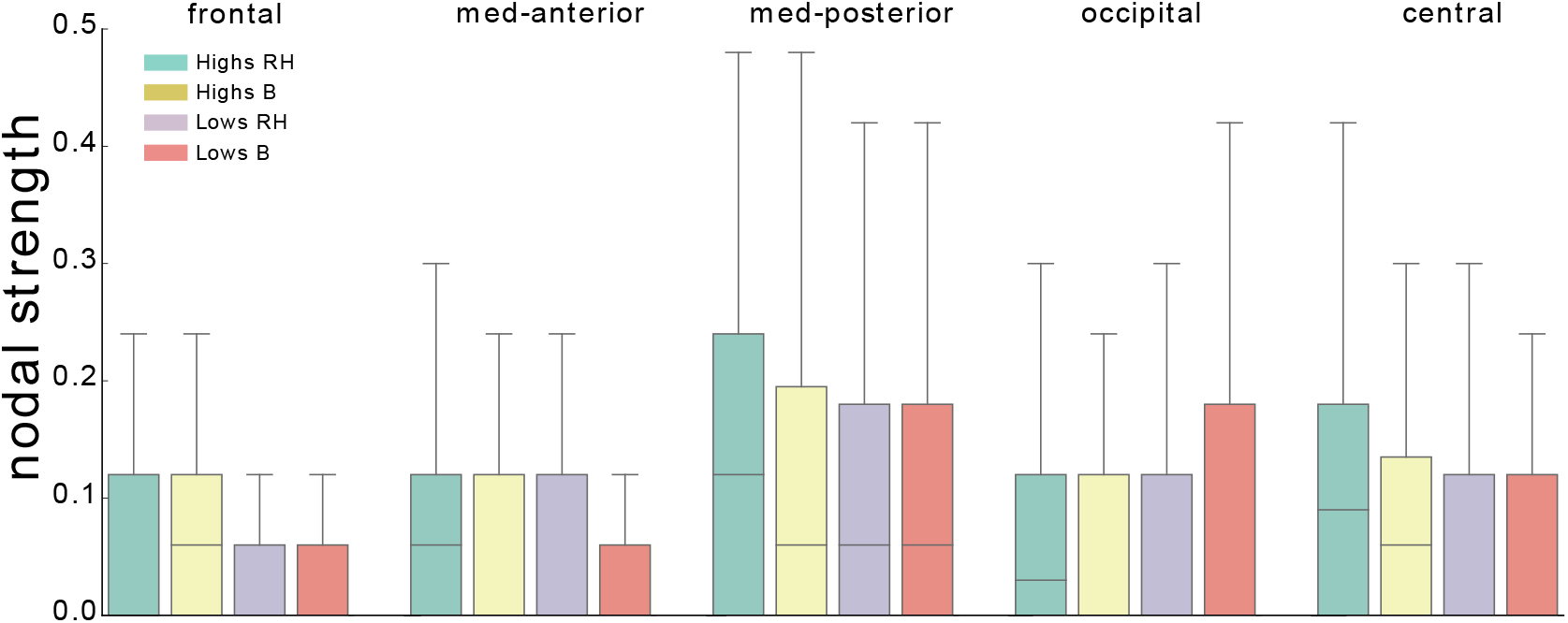
Distribution of nodal strengths. We show the distributions of nodal strengths in the scaffolds associated to the four conditions *highs*/*lows* B/RH. Nodes are grouped as follows. Frontal FP1, FP2, F8, F7, F4, F3; med-anterior: T3, FT7, T4, FC4, FT8, FC3; med-posterior: C4, C3, CP3, CP4, TP8, TP7; occipital: O1, P4, P3, PO2, O2, T6, PO1, T5; central: CZ, PZ, FZ, OZ, CPZ, FCZ.

**Table C.1:** Significant Condition effects through ANOVA test (F: Fisher test). On different EEG bands: beta1(1316 Hz), beta2 (1620 Hz), beta3 (2036 Hz) and gamma (36-45 Hz)

| Region | effect | beta1 | beta2 | beta3 | gamma |
| --- | --- | --- | --- | --- | --- |
| frontal | B<RH | F=4.050,<br>p<.051,<br>$\mu^2=.094$ | F=21.499,<br>p<.0001,<br>$\mu^2=.355$ | F=44.68,<br>p<.0001,<br>$\mu^2=.539$ | F=63.410,<br>p<.0001,<br>$\mu^2=.618$ |
| medio-anterior | B<RH | F=6.145,<br>p<.018,<br>$\mu^2=.132$ | F=30.423,<br>p<.0001,<br>$\mu^2=.438$ | F=54.514,<br>p<.0001,<br>$\mu^2=.583$ | F=69.752,<br>p<.0001,<br>$\mu^2=.641$ |
| medio-posterior | B<RH | | F=27.452,<br>p<.0001,<br>$\mu^2=.413$ | F=56.135,<br>p<.0001,<br>$\mu^2=.590$ | F=81.159,<br>p<.0001,<br>$\mu^2=.675$ |
| occipital | B<RH | | F=17.496<br>p<.0001,<br>$\mu^2=.310$ | F=48.754,<br>p<.0001,<br>$\mu^2=.556$ | F=73.985,<br>p<.0001,<br>$\mu^2=.661$ |
| midline | B<RH | | | | F=75.827,<br>p<.0001,<br>$\mu^2=.660$ |

**Table C.2:**
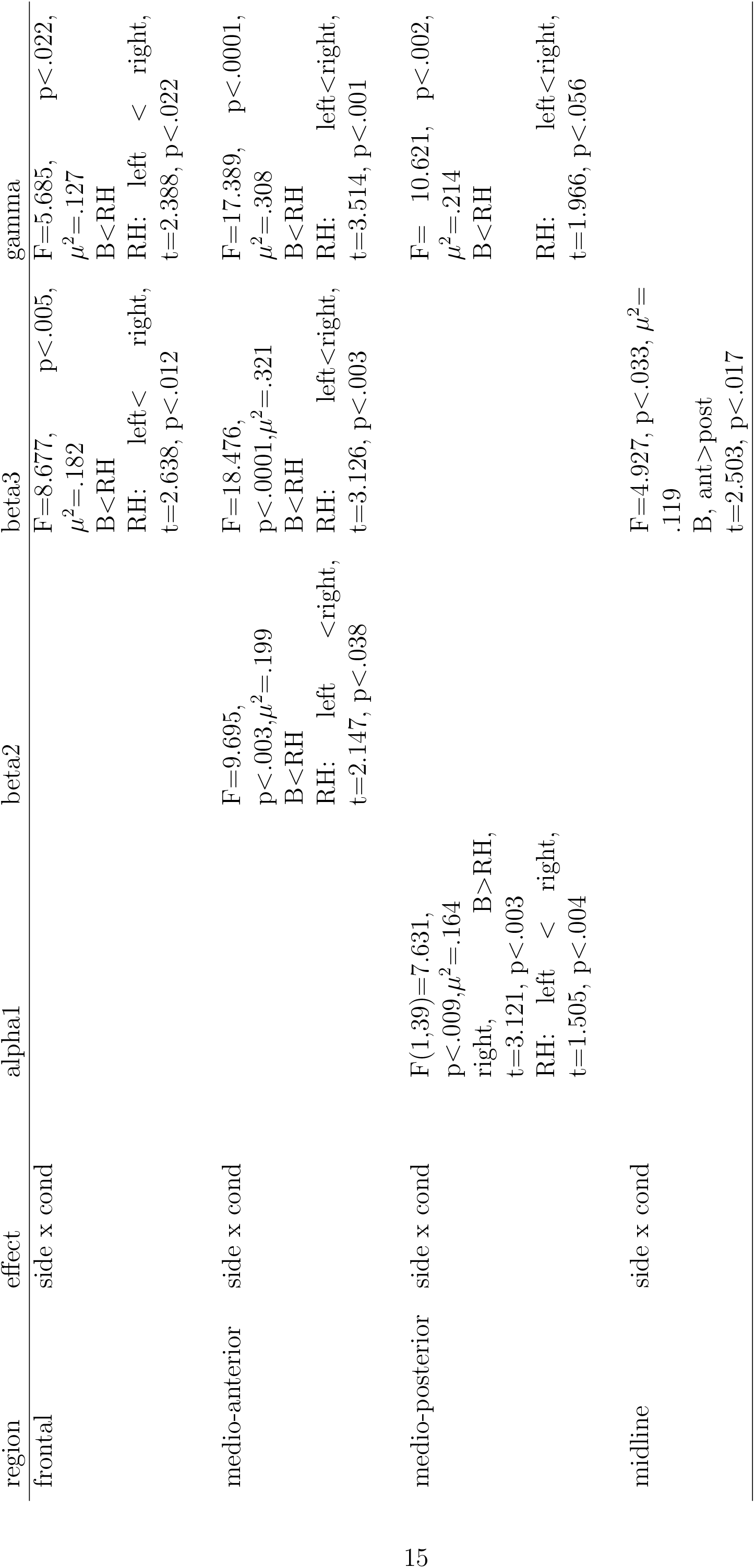
Significant condition effects through ANOVA test (F: Fisher test) on different EEG bands: beta1(1316 Hz), beta2 (1620 Hz), beta3 (2036 Hz) and gamma (36-45 Hz).

A significant interaction of Hypnotizability with Side was found for beta 2 at frontal level (*F* (1, 39) = 4.126*, p < .*049, *μ*^2^ = .096). Its decomposition revealed significant lower beta2 power on the left than on the right side in lows independently of the head position (*t*(1, 39) = 2.35, *p < .*03), whereas highs did not exhibit any asymmetry. Spectral analysis did not reveal any modulation of the EEG correlates of the rotated position of the head by hypnotizability.

### 3.2 Topological Data Analysis

Topological observables revealed significant differences between head positions and hypnotizability groups. Cycles in both groups of participants were detected that were contained in the left or right hemisphere only (pure), that were contained in the left or right hemisphere together with the involvement of some central regions (non pure) and finally that spanned regions in both left and right hemispheres. Their number, not significantly different between conditions and groups, is reported in Table C.3. However, the number of cycles not restricted to one hemisphere (non-pure + mixed) was systematically larger in both groups. Thus, ANOVA was performed on the entire frontal, medio-anterior, medio-posterior, occipital and midline region.

**Table C.3:** Classification of subjects according cycle’s composition. Numbers represent how many subjects are in each class. *pure*: all components of the cycle belong only to a one hemisphere, *non-pure*: all components of the cycle are on one hemisphere and some in central regions and *mixed*: components of the cycles belong to both hemispheres.

| group | condition | pure |  | non-pure |  | mixed | non-pure+mixed |
| --- | --- | --- | --- | --- | --- | --- | --- |
|  |  | Left | Right | Left | Right | Left-Right |  |
| *highs* | B | 9 | 8 | 12 | 8 | 20 | 40 |
|  | RH | 10 | 9 | 19 | 13 | 17 | 49 |
| *lows* | B | 13 | 10 | 15 | 11 | 18 | 44 |
|  | RH | 15 | 10 | 14 | 14 | 17 | 45 |

Significant differences between RH and B were observed in highs on right frontoparietal sites (F8 (B*>*RH, t=-3.57, p=0.002;) B*<*RH, P4 (t=2.4726, p=0.024) and in lows on left parieto-occipital sites (B*>*RH; T5 (t =2.480, p = 0.023; PO1 (t =-2.453, p=0.024), respectively.

Furthermore, as shown by the Euclidean distances computed between the vectors of nodal strength between RH and B, we find that across all subjects changes occurring in *highs* were systematically smaller than those observed in *lows* (Figure 3).

**Figure 3:**
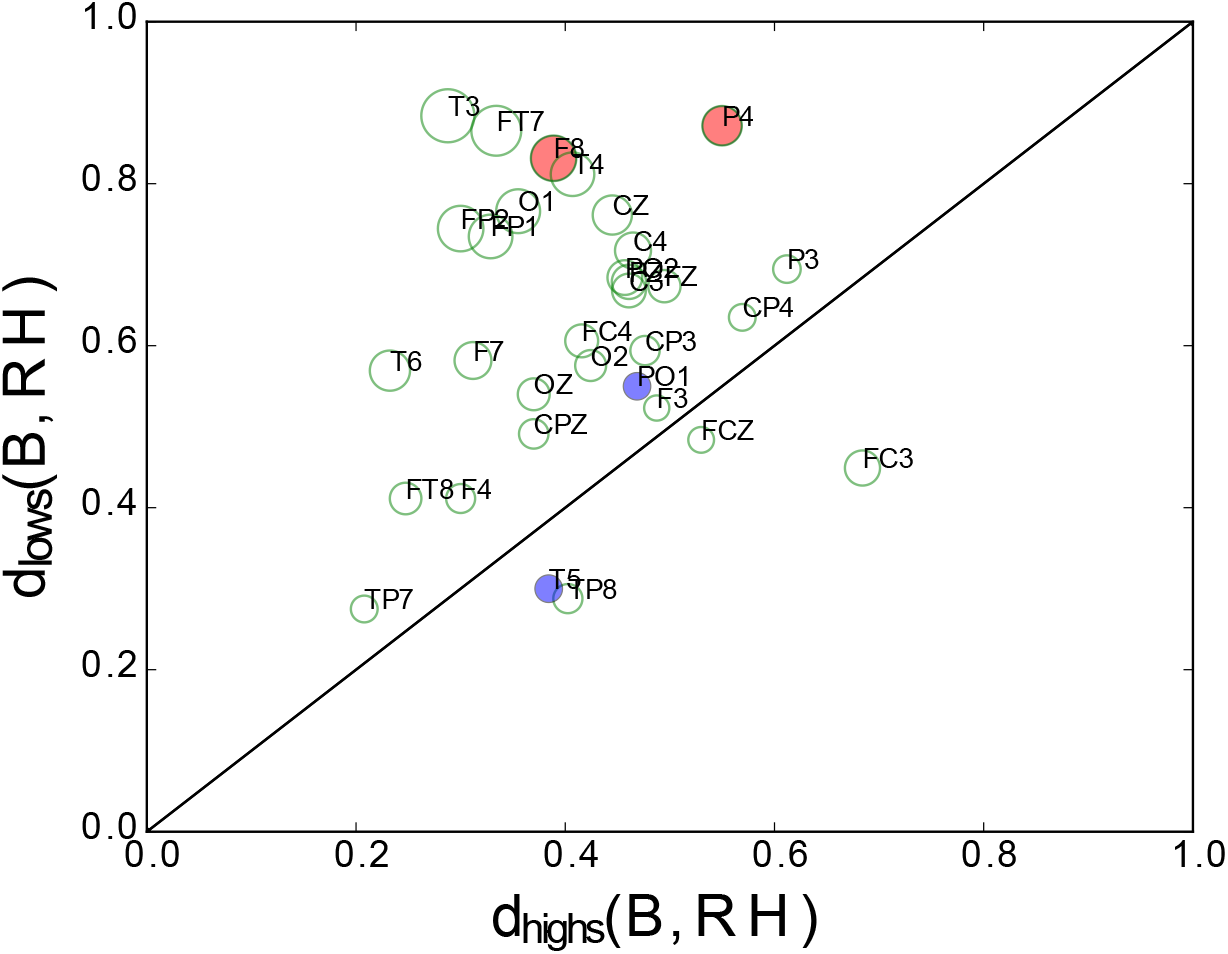
Difference between Euclidean distance of nodal strength vectors. The size of the dots is proportional to their distance from the diagonal, thus smaller points indicate smaller differences between B-RH distances. Most of the points remain in the upper diagonal part, which indicates smaller differences between RH and B in *highs*. Red and blue circles indicate significant nodal strength differences between RH and B in both groups.

Collapsing nodes by regions (Figure 3, definition of the regions are given in the captions), a significant Hypnotizability x Condition interaction was observed at occipital level (F(1,37) = 2.683, p=0.046). Its decomposition revealed only a significant difference between highs and lows during RH (t= 2.67, p=.008) in the presence of similar values in basal conditions.

## Discussion

The study provides the first report on the cortical representation of the sensorimotor context associated with the rotated position of the head. Spectral analysis captured only differences between head positions while persistence homology revealed also hypnotizability-related differences. Spectral analysis showed that the EEG power of high frequency bands increased during the maintenance of the rotated position of the head. This agrees with the beta synchronization observed during focused motor attention [Kristeva-Feige et al., 2002] and in the maintenance of postures [Gilbertson et al., 2005, Schoffelen et al., 2005], which may be accounted for by earlier observations of synchronous firing of neurons during sustained contractions [Conway et al., 1995] and slow movements of hand muscles [Salenius et al., 1997]. In contrast, alpha power decreased in the medioposterior region, in line with the inhibitory hypothesis of alpha rhythm [Palva and Palva, 2007, Palva and Palva, 2011]. Since the increases in beta and gamma power were larger on the right side (frontal and medio-anterior regions) and alpha power decreases were found only on this side (medio-posterior region), we argue that the observed EEG changes do represent the sensori-motor context associated with the rotated head posture. The asymmetric changes in the EEG power could be due to the larger proprioceptive information arising from the lengthened left neck muscles and are in line with the gamma power increases observed contralaterally to a sustained tactile stimulation [van Ede et al., 2014]. Nonetheless, in the high frequency bands the rotated head position was characterized by power increases on both sides, although larger on the right one. This may be accounted for by the two different processes characterizing our task. One was cognitive and consisted of directing attention to maintain the head rotated toward one side, the other was sensorimotor and consisted of the cortical representation of the sensori-motor asset relative to the head position. In fact, in the frontal regions lower beta (beta 1, beta2) and gamma did not exhibit any asymmetry associated with the rotated position of the head, which is consistent with the involvement of the anterior brain region in cognitive rather than sensory processes and with the bilateral representation of the cognitive processes related to movement and posture [Cremades and Pease, 2007]. Interestingly, a bilateral representation of the rotated position of the head was observed also at occipital level, where beta2, beta3 and gamma power were influenced by the head position despite the major visual competence of this region. A contribution of the occipital region may be due to cross-modal sensory activation [Heimler et al., 2015] which has been observed in several experimental protocols such as sighted adults who recruit the ventral visual cortex during tactile Braille reading [Bola et al., 2016] and, for the auditory modality, congenitally deaf subjects showing activation of the auditory cortex during tactile stimulation [Levänen et al., 1998, Poirier et al., 2005]. Nonetheless, since the occipital increases in beta and gamma were bilateral and symmetric, we hypothesize that they could be due also to a supra-modal representation of the sensory context, independent from the specific sensory modality [Bonino et al., 2015, Papale et al., 2016]. The other process that is the sensory-motor representation of the rotated head could have its correlates in the asymmetric increases in beta2, beta3 and gamma power in the medio-anterior region [Turella et al., 2016, van Ede et al., 2014]. Finally, the power increases observed on the midline sites can be related to both cognitive and sensory aspects of the task [Başar et al., 1997, Klimesch, 1999]. The scarce hypnotizability-related differences observed are in line with earlier studies of cognitive tasks characterized by more widespread changes within highs than within lows and the absence of local significant differences between highs and lows [Cavallaro et al., 2010]. With respect to spectral analysis, persistent homology provides a different perspective of the EEG changes occurring in the present study because it reveals the strength of the relation between cortical sites and can suggest the mechanisms leading to the observed spectral changes. It is more sensitive than spectral analysis to the hypnotizability-related changes in the sensori-motor context associated with the position of rotated head. The Euclidean differences between the nodal strength in RH with respect to basal conditions (Figure 3), in fact, were almost always lower in highs than in lows. Thus, on one hand the present study supports earlier findings showing larger changes in lows than in highs during both the real and imagined rotated position of the head [Ibanez-Marcelo et al., 2018]. On the other hand it reveals spatial differences and different changes between the two groups, as highs decreased their nodal strength at right fronto-parietal sites and lows at left parieto-occipital sites. In line with this observation, collapsing sites of each region a significant difference between groups during head rotation was found in the occipital region, which suggests that the maintenance of the rotated posture of the head was associated in lows with its visual representation and in highs with a preferential kinaesthetic representation, and is in line with the preferential sensory modality of imagery reported by highs and lows in earlier experiments [Santarcangelo et al., 2010].

## Limitations and Conclusions

A limitation of the study is the absence of medium hypnotizable participants. According to the many reports of gaussian distribution of hypnotizability [De Pascalis et al., 2000, Carvalho et al., 2008] they could better represent the general population. Nonetheless, bimodal distribution of hypnotizability showing a larger percentage of low-to medium hypnotizable individuals, have also been reported [Balthazard and Woody, 1989]. Thus, we think that our findings in lows are reliably referable to the general population. Another limitation is the low number of recording sites with the absence of source analysis and of electromyographic recording of neck muscles. The latter would allow to test the hypothesis of greater coherence between EEG beta and muscle activity which has been observed during the maintenance of postures [Conway et al., 1995]. Despite these limitations, this is the first study providing findings concerning the EEG representation of the integrated static vestibular/neck proprioceptive information. In addition, it indicates hypnotizability-related differences in line with earlier observations of hypnotizability-related differences in the sensori-motor domain [Santarcangelo and Scattina, 2016, Ibanez-Marcelo et al., 2018]. Finally, it shows that Topological Data Analysis is more powerful than spectral analysis in capturing such differences. In conclusion, findings support the view that hypnotizability is associated with physiological characteristics apparently unrelated to the proneness to accept suggestions [Santarcangelo and Scattina, 2016] and, thus, it may be relevant to several aspects of everyday life.

## Appendix A Mathematical formulation of homology and persistent homology

We provide here a few elementary notions required for the paper, a thorough review can be found in [Edelsbrunner and Harer, 2008] and [Hatcher, 2002]. We will limit ourselves to the following notions:

- A *clique* is a subset of vertices such that they induce a complete subgraph (*K_i_* is the full connected graph of *i* nodes). That is, every two distinct nodes in the clique are adjacent.
- A *k-simplex* is a set of *k* + 1 vertices *σ* = [*p*_0_*, …, p_k−_*_1_]. It is easy to see that it is possible to map *k*-cliques onto (*k −* 1)-simplices.
- A *simplicial complex* is a topological space constructed by simplices where simplices are points, lines, triangles, and their *n*-dimensional counterparts. We obtain a simplicial complex from a binary network by mapping *k*-cliques to (*k −* 1)-simplices.
- A *filtration* used for the computation of persistent homology consists in a family of ordered clique complexes, *X_i_* one inside the other (*• • •* ⊆ *X_i−_*_1_ ⊆ *X_i_* ⊆ *X_i_*_+1_ ⊆ *…*) obtained from the progressive thresholding of a weighted network [Petri et al., 2014].

We then defined the set of *n*-dimensional chains *C_n_*(*X*) of a simplicial complex *X* as the formal sums of *n*-simplices:

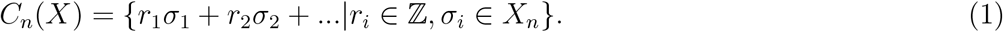

The *boundary map ∂_n_* between *n*-dimensional chains *C_n_*(*X*) to (*n −* 1)-dimensional chains *C_n−_*_1_(*X*) corresponds to the intuitive notion of boundary of a shape:

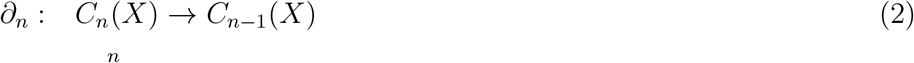

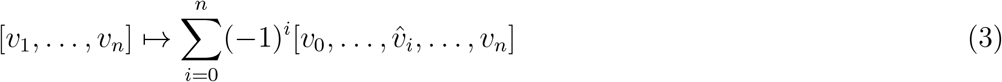

where the hat denotes the omission of the vertex. It is easy to see that *∂_n_∂_n_*_+1_ = 0 ∀*n*, that is a boundary has no boundary.

A simplicial complex *X* induces the *chain complex*, ⋯*→ C_n_*_+1_ *→ C_n_ → C_n−_*_1_ *→* ⋯*•* through boundary maps *…∂_n_*_+2_*, ∂_n_*_+1_*, ∂_n_, ∂_n−_*_1_*, …*

The *n-homology* of this complex is defined as the quotient of two vector spaces: the kernel of the map *∂_n_* quotiented by the image of the boundary map one upper dimension, *∂_n_*_+1_,

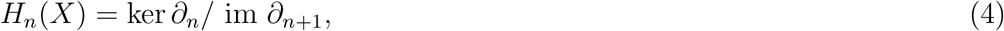

where *n* indicates the dimension of the generators in the homology group. We call the kernel ker *∂_n_* the *n*th cycle module as it usually denoted by *Z_n_*, while the image im *∂_n_* is the *n*th boundary module, denoted by *B_n_*.

For a simplicial complex *X*, a filtration is a totally ordered set of subcomplexes *X_i_* ⊂ *X*, that starts with the empty complex and ends with the complete complex, indexed by the nonnegative integers, such that:

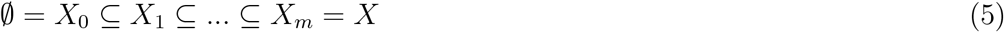

such that if *i ≤ j* then *X_i_* ⊆ *X_j_*.

In order to define *persistent homology* [Edelsbrunner and Harer, 2008, Zomorodian and Carlsson, 2005 we use superscripts to denote the index in a filtration. The *i*th simplicial complex *X_i_* in a filtration gives rise to its own chain complex 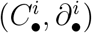 and the *k*th chain, cycle, boundary and homology modules are denoted by 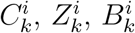 and 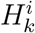, respectively.

For a positive integer *p*, *the p-persistent kth homology* module of *X_i_* is

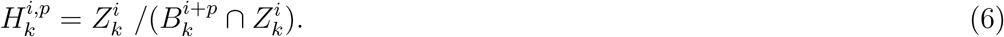

The expression for 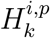 is reminding of the expression for 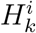, with the notable difference that it characterizes the *k*-cycles in the *X_i_* subcomplex that are not the boundary of any (*k* + 1)-chain from the larger complex *X_i_*_+*p*_, rather than those not coming from a (*k* + 1)-chain in *X_i_*. In this way 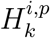 characterizes the *k*-dimensional holes in *X_i_*_+*p*_that persisted from the subcomplex *X_i_*.

The output of persistent homology can be summarized in a topological invariant called persistence diagram. A *persistence diagram* is a set of tuples (*b, d*), that describe the appearance *b* (birth step) and disappearance *d* (death step) of each hole along the filtration, with holes having longer *persistence π* = *d − b* being generally considered more relevant.

A second homological summary obtained from persistent homology is the *persistence homological scaffold*, defined for the first homology group and introduced by Petri et al. [Petri et al., 2014]. It is usually used when the original data come in the form of a network and allows to re-encode some of the information from persistent homology in a more easily interpretable network format.

It is built as follows: we consider only the case of *H*_1_; for each hole *c* we assign the shortest representative cycle and weigh its edges according to the holes’ persistence *π_c_*.

Given a graph *G*, *persistence homological scaffold* 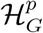, is the network composed of all the cycle *c* corresponding to generators in *H*_1_ weighted by their persistence. When an edge *e* belongs to multiple cycles *c*_0_*, c*_1_*, …, c_s_* its weight is defined as the sum of the generators persistence:

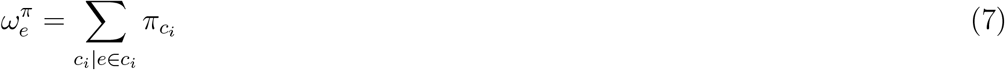

Given a weighted network *G*, with a set of nodes *n_i_* the *nodal strength*, *ns_i_*, is the sum of weights of all incoming edges to *n_i_*:

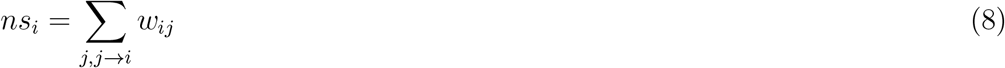

## Acknowledgements

EIM and GP were supported by the ADnD project of Compagnia San Paolo. EL was supported by the University of Pisa by the Fondi di Ateneo 2016.

## Author contributions statement

ELS designed the experiment, EIM and GP designed the analytical framework, LC and ELS conducted the experiment and pre-processed the data, EIM and GP analysed the results. All authors contributed to writing the paper and approved the final manuscript.

